# TDG regulates cell cycle progression in human neural progenitors

**DOI:** 10.1101/140319

**Authors:** I Germanguz, J Park, J Cinkornpumin, A Soloman, M Ohashi, WE Lowry

## Abstract

As cells divide, they must replicate both their DNA and generate a new set of histone proteins. The newly synthesized daughter strands and histones are unmodified and must therefore be covalently modified to allow for transmission of important epigenetic marks to daughter cells. Human pluripotent stem cells (hPSCs) display a unique cell cycle profile, and control of the cell cycle is known to be critical for their proper differentiation and survival. A major unresolved question is how hPSCs regulate their DNA methylation status through the cell cycle, namely how passive and active demethylation work to maintain a stable genome. TDG, an embryonic essential gene, has been recently implicated as a major enzyme involved in demethylation^1^. Here we present new data showing that TDG regulates cell cycle related gene expression in human neural progenitors (NPCs) derived from hPSCs and controls their capacity for neural differentiation. These observations suggest that TDG and active demethylation play an important role in hPSC cell cycle regulation and differentiation.

## Introduction

Coordinated changes to the epigenome are known to be essential for lineage specification and maintenance of cellular identity. DNA methylation and histone modifications critically contribute to epigenetic maintenance of chromatin structures and gene expression programs. DNA methylation can silence genomic regions, directly or indirectly, and play an important role during mammalian development. Loss of methylation in specific locations is associated with differentiation towards specific germ layers as binding of several transcription factors is strongly associated with specific loss of DNA methylation in one germ layer, and in many cases a reciprocal gain in the other layers^2^. However the mechanism for the lineage related site specific demethylation is not currently known. A major open question is whether this is the result of an active or passive demethylation mechanism. Promoters with low CpG content are more likely to be methylated in ESCs but demethylated and actively expressed during differentiation in a cell-type-specific manner^3^. Demethylation can occur by a passive mechanism in which the normal function of DNMT1/UHRF1 is insufficient or disrupted^4^. Alternatively, evidence for the existence of an active mechanism in which the cytosine modifications are enzymatically removed is accumulating^5^. Which of these mechanisms is responsible for demethylation changes in early human development is not currently known.

We and others have shown that standard differentiation protocols of hPSC derive an embryonic-like cell rather than a mature cell^6^. We have previously identified a group of embryonic related genes which are differentially expressed in the PSC progeny of all three lineages and in tissues of early gestation period rather than in their respective cell types of later developmental stages^6^. Among these genes we identified Thymine DNA Glycosylase (TDG), a gene that was recently implicated in active DNA demethylation (reviewed in^5^). Unlike other glycosylases, TDG is essential for embryonic viability as TDG null embryos die around E11.5-12.5 of internal hemorrhage^7,8^. The lethal phenotype is also associated with aberrant promoter methylation and imbalanced histone modifications. In addition there is evidence that levels of this enzyme are linked to progression through specific cell cycle stages. Here we describe new data showing that TDG regulates cell cycle related gene expression in human neural progenitors (NPCs) derived from hPSCs and controls their capacity for differentiation towards neurons and glia. These observations suggest that TDG and active demethylation play an important role in hPSC cell cycle regulation and differentiation.

## Results

### Expression of DNA demethylases through neural development

We originally identified TDG in a screen for genes that were consistently differently expressed between human pluripotent derivatives and their *in vivo* counterparts. This screen identified a number of genes that were persistently expressed in pluripotent derivatives and therefore suggestive of an early embryonic state, or genes that failed to be induced in pluripotent derivatives but were highly expressed in tissue derived cells. TDG was expressed significantly higher in fibroblasts, hepatocytes and neural progenitors generated from pluripotent stem cells as opposed to the same cell types derived from skin, liver and brain respectively (Fig 1A and cemagenes.com, an online repository).

**Figure 1.**
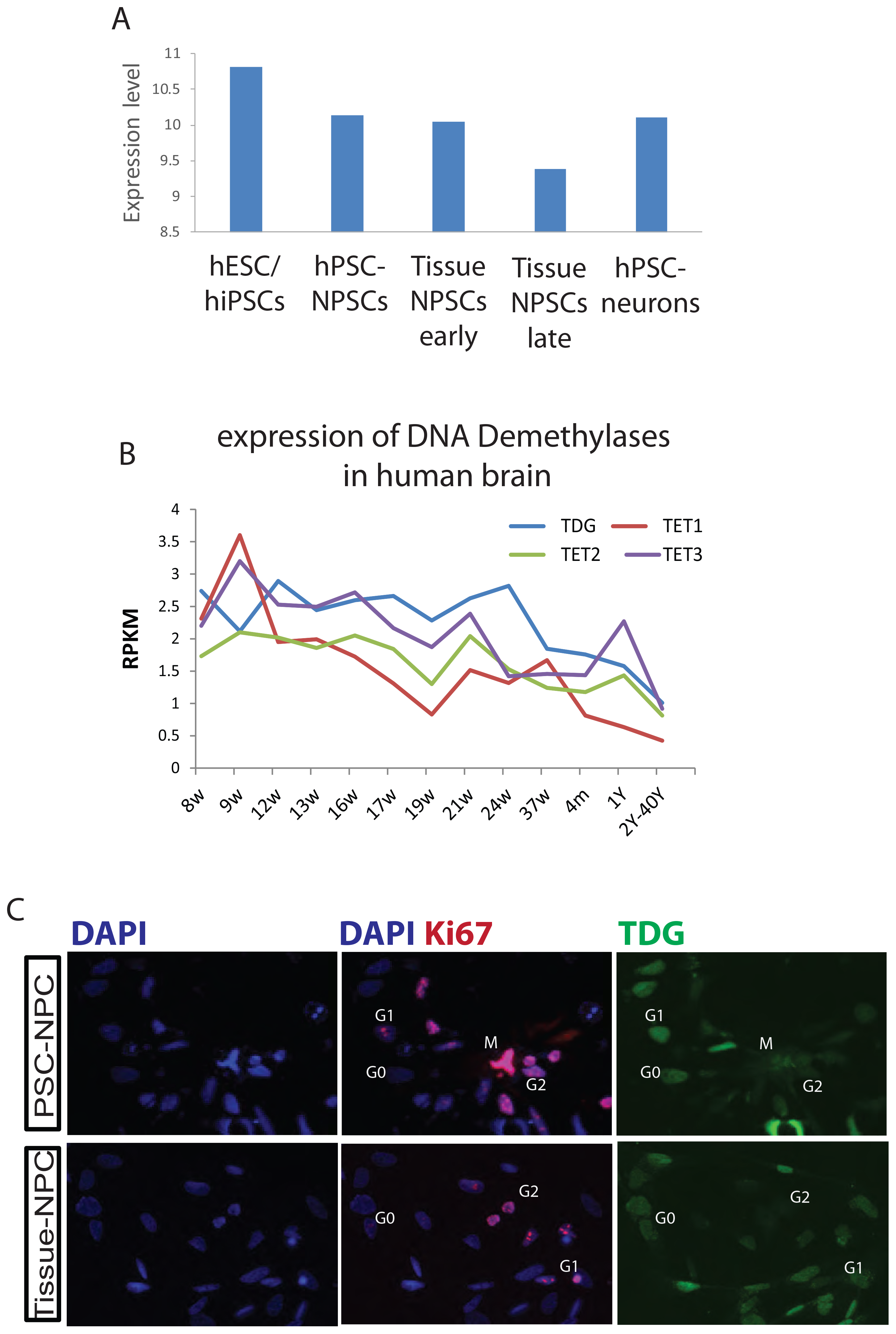
TDG is expressed during early human neuronal development. **A**, Average expression of *TDG* microarray probe in hPCS, hPSC derived NPCs and neurons and tissue derived NPCs, adapted from^12^. **B**, RNA-seq analyses for the expression of TDG, TET1,2 and 3 adopted from the Allen Institute’s Brainspan developmental transcriptome database displayed as log-scale reads per kilobase measured (log_2_ RPKM) across the developing human brain. **C**, Immunofluorescent (IF) staining for KI67 and TDG expression in NPCs derived from human pluripotent stem cells (PSC-NPCs) or human brain tissue (Tissue-NPCs). KI67 based cell cycle determination was performed according to ^9^.

The Allen Brain Atlas created by the Allen Institute provides gene expression data from various brain regions across both development and through adulthood. As shown in Fig 1B, TDG is expressed most highly in the brain *in utero*, and then falls after birth and stays low throughout adulthood. The same was true for the TET family of digoxigenases, suggesting that DNA Demethylation is primarily performed *in utero*. It is also possible that DNA Demethylation by TDG and TETs is linking to proliferation, which is known to decrease at birth relative to *in utero*. Because of this and previous data suggesting TDG could potentially regulate the cell cycle, we stained neural progenitors made from human pluripotent stem cells or derived from tissue for TDG and Ki67. The pattern of Ki67 staining and localization within the nucleus can also be used as a measure of the stage of the cell cycle. The cell cycle phase for this assay was determined based on a previously reported KI67 staining pattern^9^. TDG was previously reported to be tightly regulated during the progression of the cell cycle as its level is rapidly downregulated by ubiquitination in the S phase of the cell cycle in cellular models such as HeLa and fibroblasts and re-expressed in G2^10,11^. Here, we found a similar result, namely that TDG protein levels appear to correlate with G0/G1stages of the cell cycle (Fig 1C).

### Silencing TDG by siRNA

To investigate the role of TDG in early human development, we used siRNA mediated knockdown (KD) of TDG in neural progenitor cells (NPCs) derived from hPSC (Figure 2A). To determine whether TDG KD led to expected changes in 5-carboxylcytosine (5caC) and 5-formylcytosine (5fC) DNA residues, immunostaining for these marks was performed. As expected, silencing TDG led to increased intensity of 5caC and 5fC DNA residues, with no change in 5-Hydroxymethylcytosine (5hmC) DNA (Fig 2B).

**Figure 2.**
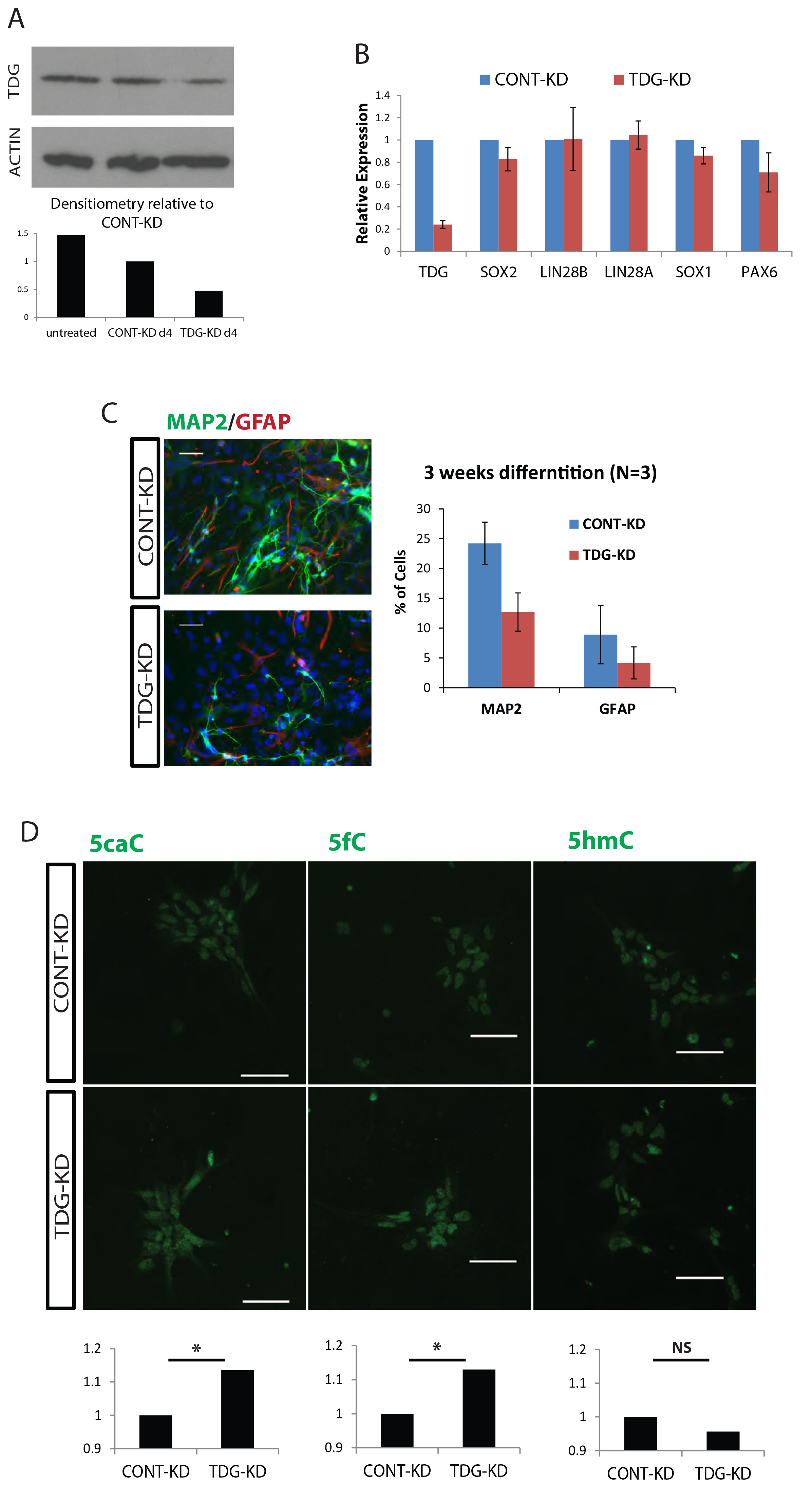
TDG downregulation in NPCs effects differentiation capacity. **A**, TDG protein expression level at day 4 post siRNA transfection, measured by Western blot (top). **B**, Expression levels of NPC markers measured by qRT-PCR, normalized against the relative levels of GAPDH and compared to CONT-KD, error bars represent standard error of the mean of 3 knockdown experiments. **C**, 4 Days post siRNA transfection, NPCs were induced to terminally differentiate by growth factor withdrawal (GFWD). Left: representative IF of 3 Weeks neural differentiation. Efficiency measured by MAP2 (neuron) /GFAP (glia). Right: quantification of n=3 separate knockdown/differentiation experiments. **D**, Top: Representative immunofluorescence of DNA methylation modifications Bottom: ImageJ quantification TDG-KD to CONT-KD ration of over 100 nucleuses across at least 5 view plains. p values were calculated with Student’s t test: *=p<0.05, ns=not significant.

RT-PCR for genes typical of the NPC state showed essentially no change in TDG KD cells (Fig 2C). 4 days post KD NPCs were induced to further differentiate using the growth factor withdrawal method (removal of self-renewal supporting growth factors EGF, bFGF^12^). Three weeks after induction of differentiation, we analyzed the percentage of MAP2/GFAP positive cells which represent the differentiation expectancy towards the neuronal/glial lineage respectfully. We found that though the neural/glial ratio remained similar, the total differentiated cell percentage was lower than in control (Figure 2D), indicating a failure to properly differentiate upon silencing of TDG. Typically, such a differentiation block would be due to aberrant differentiation or due to prolonged proliferative stimulus.

We also looked for gene expression changes following TDG-KD in NPCs by RNA-SEQ. 355 genes were differentially expressed by 1.5 fold across 3 independent experiments (Figure 3A). Using the DAVID annotation tool we classified those genes in functional groups (Figure 3B). Of the most significantly enriched functional annotations identified, we found 34 cell cycle related genes. Among those, genes which are major players in mitosis, CDK1, CDK10, Skp2 were upregulated. In contrast, other genes like CDC25B and CDKN1C (p57), which are inhibitors of cell cycle progression, were downregulated (Fig 3C).

**Figure 3.**
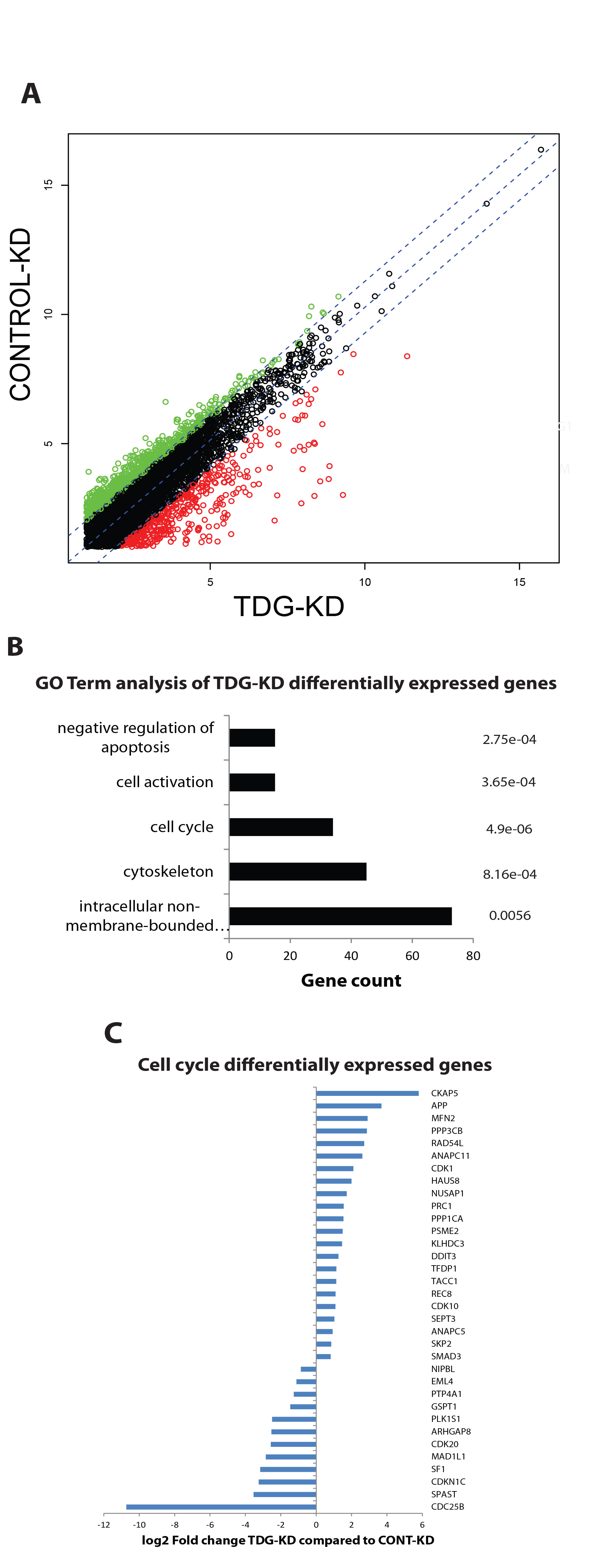
TDG-KD NPC differential gene expression analyzed by RNA-seq. **A**, Differential gene expression of n=3 siRNA knockdown experiments. Scatter plot of the group average FPKM (log2) for all genes mapped above the background cutoff, differentially expressed genes (over 1.5 fold change; p<0.05) are highlighted in red and green. **B**, Functional annotation of differentially expressed genes shows significant change in genes related to cell cycle, regulation of apoptosis and structural genes. **C**, Cell Cycle related differentially expressed list of genes and the relative fold change.

To validate that silencing of TDG by siRNA led to changes in DNA Demethylation, we performed Methylase-assisted bisulfite conversion PCR (MAB-PCR)^13^ to probe for the presence of the 5mC and 5hmC in a gene whose expression changed upon siRNA-mediated knockdown of TDG. MAB-seq takes advantage of an enzyme and bisulphite-conversion sequencing to identify the relative abundance of 5caC and 5fC nucleotides (Fig 4A). This allows for a measure of TDG activity, as TDG is known to use its glycosylase activity to finish the demethylation process to convert 5caC and 5fC to the fully demethylated state. To determine whether TDG activity can regulate the methylation and gene expression, we looked specifically at a gene whose expression was affected by siRNA-mediated silencing of TDG, EGR1, the early growth response gene which is known to be dynamically regulated by a variety of mechanisms (Fig 4B). The analyzed are was a segment of a CpG island upstream to the EGR1 TSS site, as the highest distribution of 5fC, 5caC is reported around the TSS^14^.

**Figure 4.**
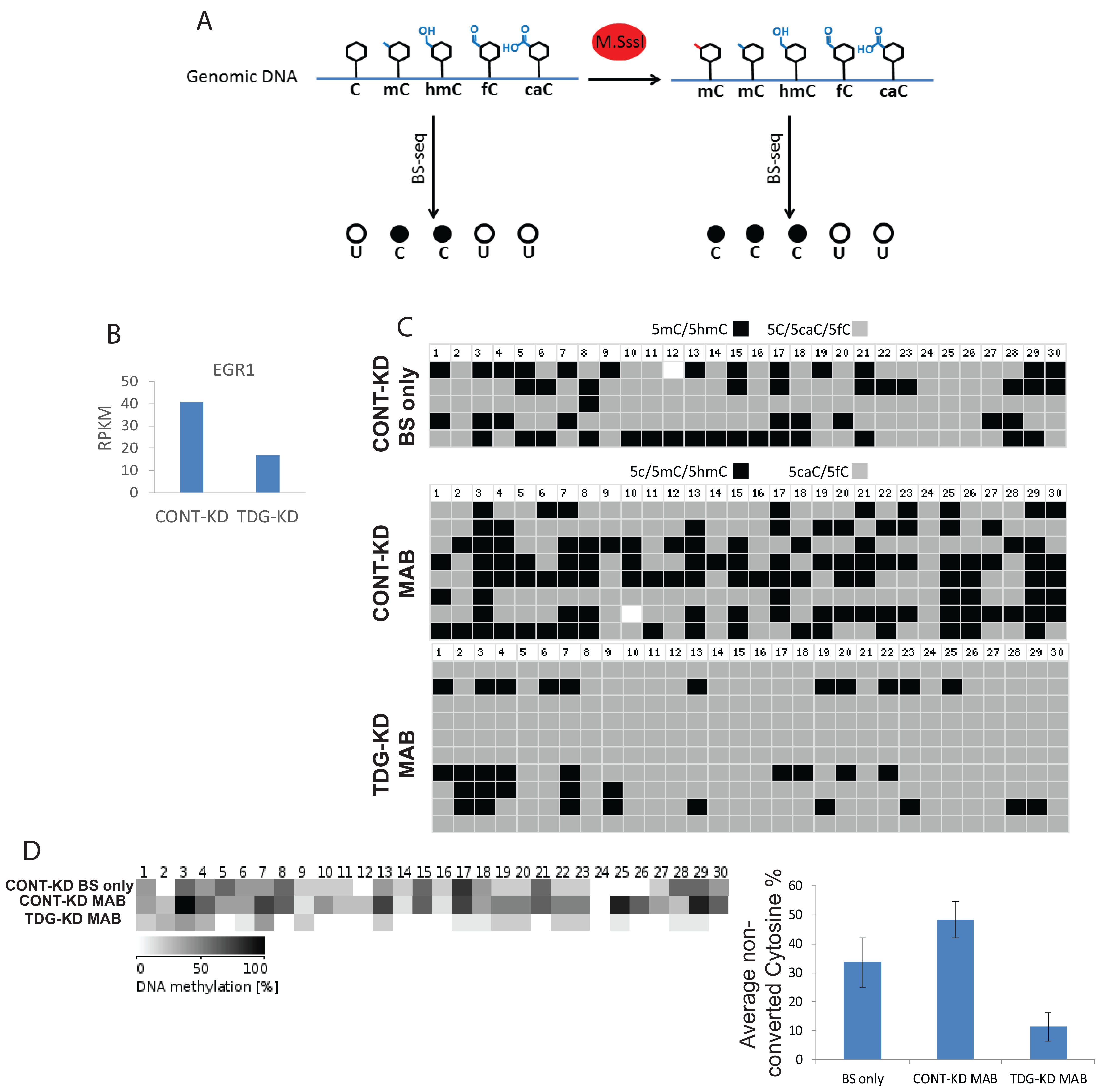
TDG downregulation results in elevation of 5caC, 5fC in the CpG island of upstream to the EGR1 TSS site. **A**, Schematic illustration of the sequencing of methylase treated compared to non-treated bisulfite converted transcripts. **B**, EGR1 expression levels are downregulated following TDG knockdown as measured by RNA-seq (described in Figure 2) **C**, Sanger sequencing of the CpG island area upstream to the EGR1 TSS following either bisulfite conversion (BS only; top) or MAB treatment (bottom two) shows higher abundance of 5cAC, 5fC in TDG deficient cells; numbers indicate CpG dinucleotide position. D, Summary visualization (left) and quantification (right) of the abundance of non-converted residues described in Figure 4C.

This analysis showed that silencing of TDG by siRNA led to a dramatic accumulation of 5caC and 5fC in a CpG island directly upstream of the start site of EGR1 transcription (Fig 4C and D). This experiment provided evidence that TDG not only regulates DNA demethylation, but also that this can influence gene expression. The proportion of genes differentially regulated by TDG-mediated DNA demethylation remains unclear until a genome-wide analysis can be performed.

We further tested whether TDG-KD in NPCs affects entrance into the cell cycle by Ki67 staining, and found that downregulating TDG resulted in a higher percentage of proliferating cells when this enzyme was knocked down (Fig 5A). Co-staining of Ki67 with TDG showed that TDG is downregulated with cell cycle progression, as higher TDG expression is observed in G0/earlyG1 cells, and downregulated with cell cycle progression (Fig 2D). We also measured cell cycle dynamics by Flow Cytometry (FACS) upon TDG silencing. This high throughput method allowed for an accurate determination of the effect of siRNA on TDG, and showed that the proportion of cells in S phase were significantly decreased, while those in G2/M were increased (Fig 5C). Taken together, we it seems clear that TDG plays a role in human pluripotent stem cell cycle regulation.

**Figure 5.**
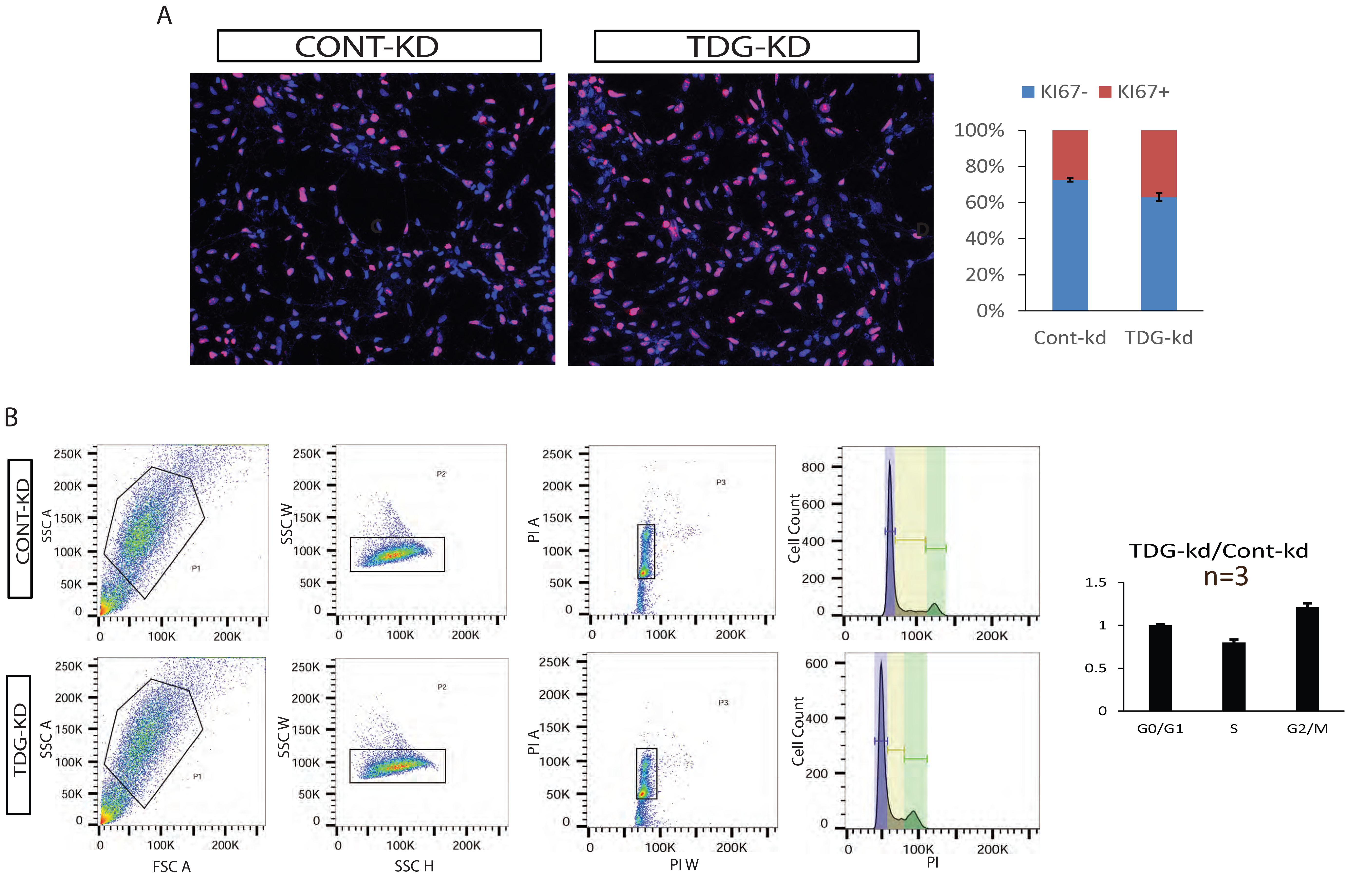
TDG deficiency entails a change in NPCs’ cell cycle. **A**, TDG downregulation results in higher fraction of cells entering mitotic cell cycle, based on KI67 positive cells staining compared to CONT-KD. Left: representative IF. Right: quantification of over 400 cells in five separate fields. **B**, TDG downregulation results with a change in cell cycle progression as G2/M increased. Left: representative flow analysis of NPC cell cycle based on DNA content staining with PI. Right: Quantification of cell cycle phase from 3 separate TDG knockdown experiments.

### Development of FUCCI model for cell cycle analyses

Despite all the analyses above, it was not clear whether silencing of TDG affects the cell cycle through its ability to regulate the terminal step of DNA demethylation. It is formally possible that the DNA glycosylation activity of TDG is used for other substrates besides methylated cytosine, for instance in DNA repair. It is also possible that another domain of TDG regulates cell cycle progression by another unknown mechanism. To attempt to link DNA demethylation by TDG to regulation of the cell cycle, we needed a system that could allow for simultaneous labeling of DNA demethylation intermediates and cell cycle markers. We generated hESCs which express the Fluorescence Ubiquitination Cell Cycle Indicator (FUCCI) transgene reporter system^15^ by lentiviral transduction. In this system cells which are in the G1 stage express Ctd1, which is conjugated to mCherry, while cells in S/G2 express Geminin which is conjugated to visible green protein mVENUS. Cells entering the DNA replication stage at the end of G1 express both markers and emit yellow light (Fig 6A). We first verified that the level of TDG is tightly regulated during cell cycle progression in hPSC as in other reported cell systems since hPSC display a unique cell cycle pattern. We found high levels of TDG in early G1 which are downregulated with cell cycle progression (Fig 6B). Interestingly, we found that 5caC and 5fC were both induced in early S phase cells, while 5hmC was reduced (Fig 6B) as was reported before ^16^. This indicates that the global state of DNA demethylation is tightly correlated to progression of the cell cycle. Furthermore, all these data on the effect of TDG on cell cycle serves to potentially explain why silencing TDG led to defective neuronal and glial specification in NPCs (Fig 2D).

**Figure 6.**
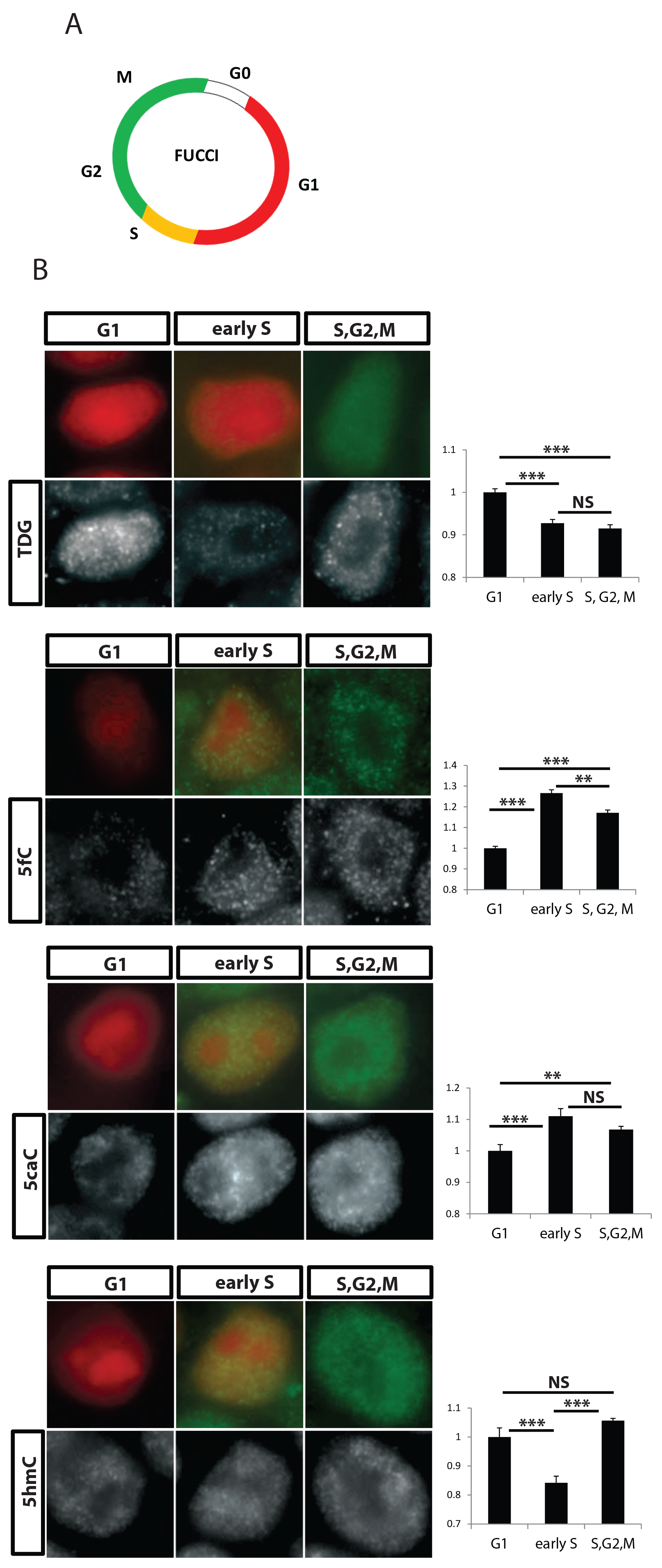
Methylation intermediate modifications level changes throughout the cell cycle. **A**, Illustration of the FUCCI cell cycle reporter system expression. **B**, Co-staining for particular cell cycle phase with antibody against either TDG or methylation intermediate modifications. Right: ImageJ quantification of over 500 cells. *p values* were calculated with Student’s t test: **=p< 0.01, ***=p<0.001, ns=not significant.

## Discussion

The data presented here confirm and extend previous findings that TDG and DNA demethylation can play a role in proper progression through the cell cycle. These results could be particularly relevant for the nervous system, where we provide evidence that TDG and TET mediated demethylation appears to diminish across development. This correlates with both proliferative rate and TDG expression, and could have important consequences to the rate of developmental progression.

The big question remaining from this work is how DNA demethylation plays a role in progression of the cell cycle. When DNA is replicated it is thought that the new daughter strand is methylated according to the hemi-methylation pattern on the sister strand by maintenance methylase (DNMT1). Less clear is what happens to portions of the genome that are hemi-methylated hydroxymethylated nucleotides. The change in proportions of 5caC and 5fC across the cell cycle could indicate that these modified nucleotides are simply erased through the action of TET and TDG enzymes, and then re-written. In this scenario, it is interesting that blocking TDG appears to promote the cell cycle rate, and could suggest that demethylation is a rate limiting step in cell cycle progression to ensure proper methylation of DNA in both daughter cells.

Because of the difference in expression levels of TDG between pluripotent and tissue derived NPCs (Fig 1A), we expected that silencing TDG would have a positive effect on the progression of developmental maturity of the NPCs. The LIN28/let-7 circuit was previously shown to be differentially regulated between NPCs born from pluripotent stem cells versus those derived from tissues, and resolution of this discrepancy was sufficient to advance the developmental maturity of NPCs in that context^17^. When the expression of TDG was brought down to a level similar to that seen in tissue derived NPCs, instead of advancing developmental maturation, the cells appeared to be unable to efficiently differentiate (Fig 2). This was presumably due to the increased rate of proliferation of the NPCs, which is known to abrogate efficient differentiation. Therefore, experimentally regulating TDG levels does not facilitate differentiation from pluripotent stem cells, as was the case with LIN28^17^. Perhaps the more interesting result from this work is that pluripotent derivatives probably need to silence TDG expression or activity at a more developmentally appropriate time point to proceed through proper development.

## Acknowledgements

We would like to acknowledge the support of various core facilities and their staff at the core facilities sponsored by the Eli and Edythe Broad Center for Regenerative Medicine (EEBCRC) including: Flow Cytometry, Genomics, and the Stem Cell Cores. This work was supported by NIH (P01GM9913) and pilot support from the BSCRC at UCLA (Rose Hills Scholar Award).

## Materials and Methods

### Tissue Culture and TDG knockdown

H9 hESCs and XFIPS2 were used in this study in accordance with the UCLA Embryonic Stem Cell Research Oversight committee. Cells were cultured in feeder free conditions on Matrigel (Corning) using mTeSR1 (Stem Cell Technologies) and passaged mechanically or with collagenase every 4-5 days. NPCs differentiation was performed as described previously^12^. Briefly, for rosette induction 80% confluent hPCS were transformed into DMEM/F12 with N2 and B27 supplements (Invitrogen), 20 ng/ml basic fibroblast growth factor (bFGF; R&D Systems), 1 μM retinoic acid (Sigma), 1 μM Sonic Hedgehog Agonist (Purmorphamine; Sigma), 10μM SB431542 (TGFβ inhibitor; Cayman) and 0.1μM LDN193189 (BMP receptor type 1 inhibitor; Cayman). Neural like rosettes were mechanically picked and expanded in DMEM/F12 supplemented with N2/B27, bFGF and 500 ng/ml epidermal growth factor (EGF; GIBCO). For further differentiation, the growth factors bFGF and EGF were with withdrawn from the media (GFW) and cultured for 3 weeks. TDG and control knockdown in NPCs was performed using a unique 27-mer siRNA duplexes (Trilencer, Origene) at a final concentration of 20 nM using Lipofectamine RNAiMAX transfection reagent (Invitrogen)

### Immunofluorescence

Immunofluorescent staining was performed using standard protocol. Briefly, cover slips were fixed with 4% PFA in PBS for 20 min, washed and then permeabilized and blocked in 10% donkey serum, 0.01% Triton in PBS for 1 hour. Primary antibodies in 5% donkey serum were incubated for 1-2 hours at room temperature following 3Xwash and incubation with conjugated secondary antibody for 1 hour in room temperature. Antibodies used include the following: rabbit anti-TDG (Atlas; HPA052263); chicken anti-GFAP (Abcam; ab4674); mouse anti-MAP2 (Abcam, ab11267), rat anti-KI67 (eBioscience; 14-5698). For methylation modifications, permeabilized cells were denatured with 2N HCl for 15 min and then neutralized with 100 mM Tris-HCl (pH 8.5) for 10 min before blocking. The following Active Motif Antibodies were used: rabbit anti 5hmC (39770); rabbit anti-5fC (61223); rabbit anti-5caC (61225).Image analysis and quantification was performed using ImageJ with the same threshold for each channel for all samples.

### RNA-Seq

Total RNA was extracted using an RNeasy Mini Kit (QIAGEN). Libraries were constructed according to manufacturer instructions (TruSeq Stranded Total RNA with Ribo-Zero; Illumina). Followed second strand PCR amplification, ~200bp sized libraries were excised from agarose gel and pooled together in 10mM concentration each. Samples were sequenced using Illumina HiSeq2000 on single-end 50-bp reads and aligned to human reference genome (Hg19) using Tophat ^18^. Processing using Cufflinks and Cuffdiff ^18^ was performed to obtain differential fragments per kilobase of transcript per million mapped reads (FPKM). Three biological replicates (i.e. 3 separate knockdown experiments in different PSC clones) were grouped together. Further analysis was performed using the cummeRbund suite ^18^. Functional annotation was performed using DAVID.

### Cell Cycle analysis

Following trypsin dissociation, knocked-down NPCs were fixed overnight in 70% ethanol in −20°. Fixed cells were then stained for half an hour at room temperature in the dark, with Propidium Iodide (PI) for a final concertation of 50 μg/ml supplemented with RNAse (final 1 μg/ml). DNA content was analyzed on BD-Biosciences LSR-II flow cytometer and cell cycle phases were determined using the FlowJo cell cycle module.

### MAB-PCR

Genomic DNA was extracted using the DNeasy Blood & Tissue Kit (Qiagen). 1 μg genomic DNA was treated by M.SssI (New England Biolabs) in a 50 μl reaction for three rounds. For each round DNA was incubated with 4 U of M.SssI CpG methyltransferase (NEB), supplemented with 160 mM final S-Adenosyl methionine for 3 hours at 37°C. At the end of each round DNA was cleaned using phenol/chloroform extraction. Bisulfite conversion was performed using the EpiTect Bisulfite Kit (QIAGEN) and then selected loci was PCR amplified with KAPA HiFi Hotstart Uracil+ DNA polymerase (KAPABiosystems). Resulting PCR product was cloned into the TOPO-Blunt (Invitrogen) vector, and Sanger sequenced. Analysis and visualization of sequence reads was done using the online BISMA tool (http://services.ibc.unistuttgart.de/BDPC/BISMA/)

### FUCCI cell lines generation

The FUCCI reporter lentiviral plasmids, pCSII-EF-mCherry-hCdt1(30/120) and pCSII-EF-mVenus-hGeminin(1/110) were a generous gift of Dr. Atsushi Miyawaki (RIKEN Brain Science Institute, Saitama, Japan). Lentiviral virions were generated in 293T cells using stranded protocols followed by concentration with Amicon Ultra-15 centrifugal units (100K; Millipore). hPSC were single celled 24h prior to infection and supplemented with Rho-associated kinase (ROCK) inhibitor Y27632 (Stemgent). Cells were first infected with one reporter followed by FACS sorting to insure that all cells are infected and then infected with the second reporter and FACS sorted again.

